# Expansion and transformation of the minor spliceosomal system in the slime mold *Physarum polycephalum*

**DOI:** 10.1101/2020.10.12.336362

**Authors:** Graham E. Larue, Marek Eliáš, Scott W. Roy

## Abstract

Spliceosomal introns interrupt nuclear genes and are removed from RNA transcripts (“spliced”) by machinery called spliceosomes. While the vast majority of spliceosomal introns are removed by the so-called major spliceosome, diverse eukaryotes also contain a mysterious second form, the minor spliceosome, and associated introns [1–3]. In all characterized species, minor introns are distinguished by several features, including being rare in the genome (∼0.5% of all introns) [4–6], containing extended evolutionary-conserved splicing sites [4,5,7,8], being generally ancient [9,10] and being inefficiently spliced [11–13]. Here, we report a remarkable exception in the slime mold *Physarum polycephalum*. The *P. polycephalum* genome contains > 20,000 minor introns—25 times more than any other species—with transformed splicing signals that have co-evolved with the spliceosome due to massive gain of efficiently spliced minor introns. These results reveal an unappreciated dynamism of minor spliceosomal introns and spliceosomal introns in general.

---

Over the nearly two decades since the surprising discovery of the existence of the minor spliceosomal system [1,2], minor (U12-type) introns have consistently been defined by a number of hallmark characteristics distinct from their major (U2-type) counterparts. First, in all lineages examined U12-type introns are either rare or absent, ranging from ∼700 (0.36% of all introns) in humans [5,6,8] to 19 (0.05%) in fruitflies [14] to complete absence in diverse lineages [4,15].

Second, U12-type introns show distinct extended splicing motifs at the 5′ splice site (5′SS) ([G/A]TATCCTT) and branchpoint sequence (BPS) (TTCCTT[G/A]AC, ≤ ∼45 bases from the 3′ splice site (3′SS)) which exactly basepair with complementary stretches of core non-coding RNAs in the splicing machinery [3,16,17]. Third, U12-type introns are typically ancient (e.g., 94% of human U12-type introns are conserved as U12-type in chicken [7]), implying low rates of U12-type intron creation through evolution [4,7,9,18]. Finally, U12-type introns show slow rates of splicing, suggesting inherently low efficiency of the U12 spliceosomal reaction [11–13,19].

## Methods

### RNA-seq based reannotation of the *Physarum polycephalum* genome

We downloaded the *P. polycephalum* genome assembly and annotation from http://www.physarum-blast.ovgu.de/, and RNA-seq for *P. polycephalum* from NCBI’s SRA database (accession numbers DRR047256, ERR089824-ERR089827, and ERR557103-ERR557120). To reannotate the genome, we combined *de novo* and reference-based approaches. First, we generated a *de novo* transcriptome from the aggregate RNA-seq data using Trinity [20] (v2.5.1). We also separately mapped the reads to the genome using HISAT2 [21] (v2.1.0), allowing for non-canonical splice sites (--pen-noncansplice 0), followed by StringTie [22] (v1.3.3) to incorporate the mapped reads with the existing annotations and generate additional putative transcript structures.

Coding-sequence annotations for the assembled transcripts, informed by additional homology information from the SwissProt [23] protein database, were generated using TransDecoder [24] (v5.0.2), and further refined with the *de novo* transcriptome via PASA [25] (v2.2.0). In addition, an AUGUSTUS [26] (v3.3) annotation was generated from the mapped reads using BRAKER1 [27] (v2.1.0) explicitly allowing for AT-AC splice boundaries (-- allow_hinted_splicesites=atac). Lastly, the AUGUSTUS- and StringTie-based gene predictions were merged using gffcompare [28] (v0.10.5), and processed again using TransDecoder. To gauge the quality of our annotations versus those previously available, we performed a BUSCO [29] (v3.0.1) analysis against conserved eukaryotic genes; the previous annotations contained matches to 60.1% of eukaryotic BUSCO groups (54.5% single-copy; 27.1% fragmented; 12.8% missing); our annotation increased this percentage to 73.3% (64.4% single-copy; 18.5% fragmented; 8.2% missing).

### Classification of intron types in *P. polycephalum*

All annotated intron sequences from our improved *P. polycephalum* genome annotation were collected and analyzed using a modified version of intronIC [8]. Briefly, we first obtained high-confidence sets of U12- and U2-type *P. polycephalum* introns, as follows. High-confidence U2-type introns were defined as introns conserved as U2-type in at least three other species. Due to the low evolutionary conservation of putative *P. polycephalum* U12-type introns, the confident U12-type intron set was assembled by combining U12-type introns conserved as U12-type in one or more species, introns with perfect 5′SS motifs ([GA]TATCCTT) interrupting coding sequences in regions of good alignment to orthologs in one or more species, introns with near-perfect 5′SS motifs in addition to the TTTGA BPS motif 10-12 bp upstream of the 3′SS, and AT-AC introns (less likely to be false positives) with strong 5′SS consensus motifs in conserved eukaryotic genes (defined as representing a BUSCO match).

Sub-sequences of each intron corresponding to the 5′SS (from −3 to +8, where +1 is the first intronic base), 3′SS (from −5 to +4, where −1 is the last intronic base) and all 12mers within the branchpoint region (−45 to −5 where −1 is the last intronic base) were scored against position-weight matrices (PWMs) derived from the sets of high-confidence *P. polycephalum* U2- and U12-type introns to obtain U12/U2 log ratio scores for each motif. These log ratios were normalized to z-scores for each motif (5′SS and BPS), and were then used to construct two-dimensional vector representations of each intron’s score. In addition, to account for the narrow window of occurrence of the non-canonical TTTGA BPS, intronIC was modified to weight the branchpoint scores of introns whose BPS adenosines were found within the range [−12, −10] of the 3′SS, with the additional weight equal to the frequency of occurrence of the BPS adenosine at the same position within confident U12-type introns. Furthermore, unless explicitly stated otherwise, we used a more conservative U12-type probability score of 95% for classifying introns in *P. polycephalum*. The prominently-separated “cloud” in the upper-right of Figure 1D is composed mainly of AT-AC U12-type introns, whose 5′SS scores are more distinct than U12-type introns with other splice boundaries.

**Fig. 1.**
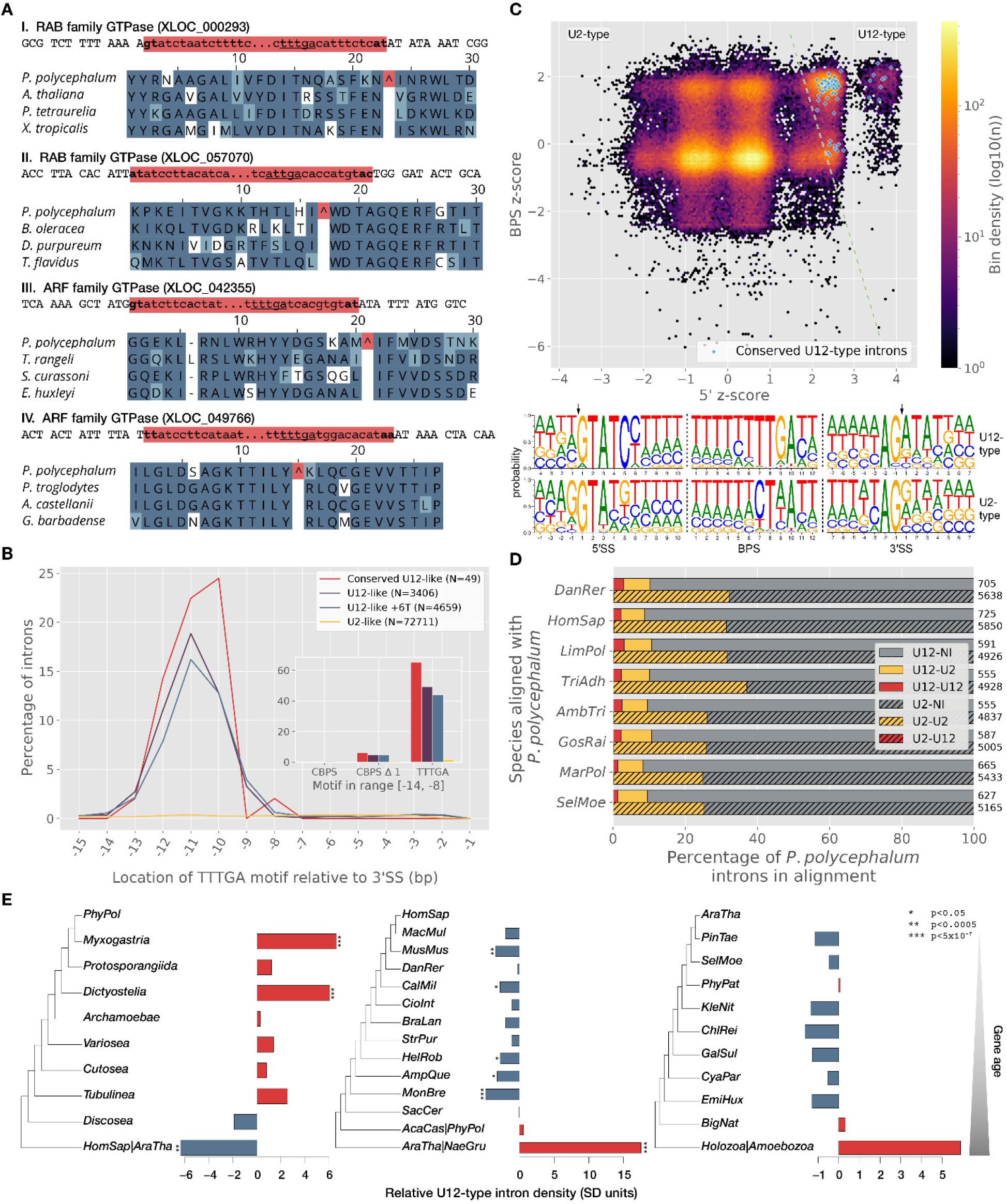
Evidence for massive gain of U12-type introns in *Physarum polycephalum*. **(A)** Canonical and non-canonical U12-like introns in conserved *P. polycephalum* GTPase genes. “^” indicates intron position. Lowercase red indicates intron sequence, with terminal dinucleotides in bold and putative BPS motifs underlined. **(B)** Presence of BPS motif in various groups of *P. polycephalum* introns. (Main) Occurrence of TTTG<u>A</u> motif as a function of number of nucleotides upstream of the 3′SS), for U12-like ([AG]TATCCTT-A[CG] or [AG]TATC<u>T</u>TT-A[CG] SS’s for “U12-like” and “U12-like +6T”, respectively), U2-like (GTNNG-AG), and conserved U12-like ([AG]TATC[CT]-NN and conserved as a U12-type intron in another species). (Inset) The same data as a cumulative bar plot for positions −14 through −8. **(C)** Conservation status of *P. polycephalum* introns in other species, showing substantially lower U12-than U2-type conservation. For each species, the pair of bars shows the fractions of *P. polycephalum* introns of each intron type (U12-type, unhashed; U2-type, hashed) that are conserved as either U12-type (red) or U2-type (yellow) introns, or not conserved (gray). Total numbers of *P. polycephalum* introns assessed are given at right. **(D)** Intron type classification and associated motifs. The main plot shows BPS-vs-5′SS log-ratio z-scores for all *P. polycephalum* introns, with conserved U12-type introns highlighted in blue. The dashed green line indicates the approximate U2-U12 score boundary (Methods, Figure S3). Below the scatter plot are sequence logos showing motif differences between the two groups (20899 U12-type, top; 154299 U2-type, bottom). **(E)** Comparison of U12-type intron density (fraction of introns that are U12-type) in genes of different age categories for *P. polycephalum* (*PhyPol*), *Homo sapiens* (*HomSap*) and *Arabidopsis thaliana* (*AraTha*), relative to expectation (blue/red = below/above expectation). U12-type intron densities in *P. polycephalum* are significantly overrepresented in newer genes, in contrast to the pattern seen in both human and *Arabidopsis*. For full species names, see Table S1.

### Identification of homologous sequences and conserved intron positions

Genomes and annotations for all additional species were downloaded from various online resources (Table S1), and in cases where sufficient RNA-seq was available and we suspected that U12-type introns had been systematically suppressed (e.g. zero or very few AT-AC introns annotated), we performed RNA-seq based annotation updates using Trinity and PASA [20,24]. For each genome, annotated coding sequences were extracted and translated via a custom Python script (https://github.com/glarue/cdseq). Annotated intron sequences were collected and scored using intronIC [8] with default settings. Under these settings, only introns defined by CDS features from the longest isoform of each gene were included, and introns with U12-type probability scores > 90% (or > 95% for *P. polycephalum*) were classified as U12-type. Furthermore, introns shorter than 30 nt and/or introns with ambiguous (“N”) characters within scored motif regions were excluded.

Between *P. polycephalum* and each other species (or, in the case of paralogs, itself), we performed pairwise reciprocal BLASTP [30,31] (v2.6.0+) searches (E-value cutoff of 10^−10^), and parsed the results to retrieve reciprocal best-hit pairs (defined by bitscore) using a custom Python script (https://github.com/glarue/reciprologs). Pairs of homologous sequences were globally aligned at the protein level using ClustalW [32] (v2.1), and introns occurring at the same position in regions of good local alignment (≥4/10 shared amino acid residues on both sides of the intron) were considered to be conserved (based on the approach in [33]).

### Calculation of dS values between paralogs

We identified 8267 pairs of paralogs in *P. polycephalum* using the same approach as for other homologs. Each pair sharing at least one intron position was globally aligned at the protein level using Clustal Omega [34] (v1.2.4), and then back-translated to the original nucleotide sequence using a custom Python script. Maximum likelihood dS values for each aligned sequence pair were computed using PAML [35] (v4.9e) (runmode = −2, seqtype = 1, model = 0), with dS values greater than 3 treated as equal to 3 in subsequent analyses (as dS values > 3 are not meaningfully differentiable in this context) (Figure S3).

### Relative gene ages

For the three focal species (FS) *P. polycephalum*, human and *Arabidopsis thaliana*, sets of node-defining species (NDS) were selected to represent a range of evolutionary distances from the FS based on established phylogenetic relationships. In the case of *P. polycephalum*, we used data from Kang et al. [36] and their amoebozoa phylogeny; for the other two FS, we downloaded the NDS genomes and annotation files from the publicly-available resources Ensembl, JGI and NCBI (Table S1). We then performed one-way BLASTP (v2.8.0+) searches (E-value cutoff 10^−10^) of each FS transcriptome against the transcriptomes of its NDS set to establish an oldest node for each gene, defined as the ancestral node of the FS and the most-distantly-related NDS where one or more BLASTP hits to the gene were found. For example, a human gene would be assigned to the human-*Danio rerio* ancestral node if a BLASTP hit to the gene was found in *Danio rerio* (and optionally any more closely-related NDS) but not in any other more distantly-related NDS.

Once gene ages were assigned, for each FS we examined the difference of the observed and expected number of U12-type introns at each node using an expected value based on the aggregate density of U12-type introns in all other nodes, and scaled the observed-minus-expected value by dividing by the node’s expected standard deviation (SD). For a given node, if *n* is the total number of introns in the node and *p* is the expected frequency of U12-type introns at the node based on combined frequencies from the other nodes, then per the binomial theorem 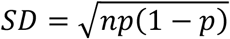. The significance of the observed numbers of U12-type introns at each node was calculated with a Fisher’s exact test (scipy v1.5.2 [37]), and *p*-values were corrected for multiple-testing using the Holm step-down method as implemented in the Python library statsmodels [38] (v0.11.1).

### Estimation of intron splicing efficiency and retention

For each annotated intron defined by CDS features from the longest isoform of each gene, splice junctions for the spliced (5′ exon + 3′ exon) and retained (5′ exon + intron, intron + 3′ exon) structures were created in silico using a custom Python script. RNA-seq reads (accession numbers listed in the reannotation section) were then mapped in single-end mode to the junction constructs using Bowtie v1.2.2 [39] with parameter −m 1 to exclude multiply-mapped reads.

Reads overlapping a junction by ≥5 nt were counted and corrected by the number of mappable positions on the associated junction construct. For each RNA-seq dataset, introns with no reads supporting the spliced form were excluded from the analysis, as were introns with no junctions supported by at least 10 reads. For all other introns, efficiency was calculated as the ratio of splice-supporting read coverage (*Cs*) over the total read coverage, which is just *Cs* plus the average of the retention-supporting read coverage (*Cr*), expressed as a percentage, i.e 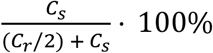. Each intron’s splicing efficiency was then computed as either the average across all RNA-seq samples containing mapped reads (for the bulk analysis), or using per-sample read counts (for the comparisons of neighboring introns from the same transcript).

To help validate our splicing efficiency results, we also employed an established method to evaluate intron retention using the same RNA-seq data. We obtained intron retention values for all annotated *P. polycephalum* introns with IRFinder [40] (v1.3.0), which produced an equivalent (inverted) pattern to our splicing efficiency metric (Figures S7-10). Introns with IRFinder warnings of “LowSplicing” and “LowCover” were excluded.

### U12-type introns in *P. polycephalum* paralogs and non-canonical U12-type introns

Introns conserved across *P. polycephalum* paralogs were identified as described for homologous introns. We then examined all intron positions conserved between paralogs, and tabulated the intron types at each position. To determine the relative likelihood of a given U12-type intron being conserved as U12-type across paralogs, we calculated the relative probability of an intron *B* being U12-type conditioned on its paralogous intron *A* being U12-type or U2-type as 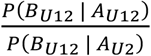, which results in a likelihood fold-increase of 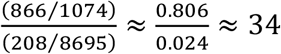. This value is most likely conservative, as decreasing the stringency of U12-type classification results in a further increase in the relative likelihood.

To avoid inclusion of spurious intron annotations representing artifacts of the RT-PCR process (“RTfacts”, [41]) in our non-canonical intron analysis, we used a fairly simple heuristic to detect unexpectedly high similarities between extended sequences around the 5′ and 3′ splice sites. For each intron, we considered regions of 24 bp centered around the 5′ and 3′ splice sites (12 bp from the exon and 12 bp from the intron in each case) and used a 12 bp sliding window to compare every 5′SS 12mer against every 3′SS 12mer. For each 12mer pair, we defined their pairwise similarity *s* as 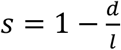, where *d* is the Hamming distance between the two strings and *l* is their length in bp (i.e. 12), and treated the highest value found as the overall similarity score. Introns with similarity scores >= 0.916 (corresponding to one mismatch between the pair of splice site 12mers) were considered possible RTfacts and were excluded (n = 1,624, 0.93% of 175,198 total introns).

In our survey of non-canonical introns in *P. polycephalum*, we took advantage of the greater number of conserved U12-type intron positions within paralogs (versus with other species) to gauge support for non-canonical U12-type intron boundaries present in our annotations. Of the non-canonical U12-type introns found in regions of good alignment between paralogs, 66% (42/63) contained the U12-type BPS motif 9-12 bp upstream of the 3′SS; in the smaller set conserved as introns between paralogs, the same motif was present in 73% (22/30). The BPS motif enrichment within these introns supports their identity as genuine non-canonical U12-type, and the distribution of the most common boundaries found within paralogs is consistent with the broader set of non-canonical U12-type introns (Figure S6B).

### Relative expression of snRNPs

Orthologs for components of the major and minor spliceosome (major: SF3a120/ SAP114, SF3a60/SAP61, U1-70K, U1 A, U2 A’; minor: U11/U12 20K, 25K, 25K, 31K, 35K, and 65K) were identified via reciprocal BLASTP searches (as described in the section on ortholog identification) using the components’ annotated human transcripts as queries (Table S2). For each species, a series of RNA-seq samples (curated by size and wild-type status) were aligned to the coding sequences of all available components using HISAT2, and the output processed with StringTie using the “-A” option to obtain per-transcript TPM values. Mean per-species TPM values across all RNA-seq samples for the U12- and U2-type components were then compared to calculate the U12/U2 expression ratios. An ANOVA test was performed on the aggregate group of ratios (*p*-value 8.8×10^−13^), followed by pairwise two-tailed Mann-Whitney U tests between all combinations of ratios. The difference between *P. polycephalum* and every other species was significant at p < 0.05 after multiple-testing correction using the Holm step-down method as implemented in the Python library statsmodels v0.11.1 [38].

## Results

### U12-type intron enrichment in *Physarum*

During manual annotation of GTPase genes in the genome of the slime mold *Physarum polycephalum*, we observed several introns lacking typical GT/C-AG boundaries, including both AT-AC and non-canonical introns (i.e., neither G[T/C]-AG nor AT-AC; Fig. 1A). Most of these atypical introns also contained extended U12-like 5′SS motifs ([G/A]TATC[C/T]TTT), consistent with previous evidence of U12 splicing in this species [15,42]. However, genome-wide analysis of the current *P. polycephalum* genome annotation revealed that all annotated introns have GY-AG boundaries, a pattern suggesting non-GY-AG introns may have been discarded by the annotation pipeline [26,43]. Indeed, an RNA-seq based genome reannotation using standard pipelines (Methods) allowing for non-GY-AG introns improved overall annotation quality (73.3% versus 60.1% BUSCO [29] sets present), and revealed a large number of previously unannotated introns, including a substantial number of introns with AT-AC splice boundaries (1,830 AT-AC, 54,816 GY-AG).

Our updated *P. polycephalum* annotation contains 3,648 introns with perfect matches to the canonical U12-type 5′SS motif (3021 with GTATCCTT, 627 with ATATCCTT). In contrast, far fewer introns exhibit a classic U12-type BPS motif (561 with CCTT[G/A]AC present in the last 45 bases out of all introns, and only 20 of the 3,648 introns with perfect U12-type 5′SS motifs), and standard position weight matrix (PWM) methods (following [6,8,42,44] failed to clearly identify U12-type introns (Methods and Figure S1). Lack of classic U12-type branchpoints were confirmed for a subset of conserved U12-type introns (those with U12-like 5′SS motifs found at positions that match those of U12-type introns in other species (Methods)). Instead, we noted the motif TTTGA falling within a short region near the 3′SS (terminal A 9-12 bp upstream of splice site), a feature also common in the manually identified non-GY-AG introns (Fig. 1A). Genome-wide analysis of the 5′SS and TTTGA motifs showed a clear correspondence: TTTGA motifs are present 9-12 bp upstream of the 3′SS in 59% (41/70) of conserved U12-type introns, as well as 42% (3,107/7,462) of GTATCYTT-AG introns and 67% (417/625) of ATATCYTT-AC introns, but only 6% (10,313/167,111) of other introns (Fig. 1B). Consistent with a role in splicing, among introns with U12-like 5′ splice sites, introns containing the TTTGA motif had lower average retention than those without it (Figure S2).

Combining this position-specific atypical branchpoint motif with species-specific splice site motifs under a support-vector machine and PWM strategy [8] (Methods) led to a clearer separation of putative U12- and U2-type introns (Figs. 1C, S3). Using a conservative criterion, we identified 20,899 putative U12-type introns in *P. polycephalum*, 25 times more than previously observed in any species. The true U12-type nature of these introns was further supported by two additional findings. First, comparisons of 8,267 pairs of *P. polycephalum* paralogs showed strong conservation of U12-type character: among intron positions shared between paralogs, an intron was 34-45 times more likely to be predicted to be U12-type if its paralogous intron was predicted to be U12-type (Methods, Figure S4). Second, putative *P. polycephalum* U12-type introns as a group are strongly biased away from phase 0 (26% compared with 39% for U2-type introns; phase is not a component of the scoring process), consistent with the phase bias observed in other species (Figure S5) [10,45].

### Evolution of *Physarum* U12-type introns

Interestingly, very few U12-type intron positions in conserved coding regions are shared with distantly-related species (e.g., only 9% of *P. polycephalum* U12-type introns found as either U2- or U12-type introns in humans, far fewer than for U2-type intron positions (31%); Fig. 1D, see Table S1 for species abbreviations), indicating either massive U12-type intron gain in *P. polycephalum* or equally massive loss in other species. There is, however, no evidence for widespread loss of U12-type introns in other species, and previous results have attested to significant U12-type intron conservation across long evolutionary distances [7,8,46]. Indeed, among U12-type introns conserved between *P. polycephalum* and plants and/or animals (i.e. ancestral U12-type introns), 63% are retained as either U2- or U12-type in the the variosean amoeba *Protostelium aurantium*, and 70% are similarly retained in the discosean *Acanthamoeba castellanii* (Figure S6). That *P. polycephalum* has recently gained many U12-type introns is also supported by the fact that putatively recently-evolved *P. polycephalum* genes (i.e., those lacking homology to genes outside of closely-related species) show substantial U12-type intron densities (Fig. 1E). This finding is not expected from retention of ancestral U12-type introns and is in clear contrast to the low U12-type intron densities in young human and plant genes (Fig. 1E).

### Features of the U12 system in *Physarum*

Analysis of the highly expanded U12 spliceosomal system in *P. polycephalum* revealed a variety of other surprising characteristics. In contrast to the remarkable consistency of splice sites in most eukaryotic genomes (e.g., 99.85% GY-AG or AT-AC in humans), we found many noncanonical introns in *P. polycephalum* (Methods). After filtering for likely reverse transcriptase artifacts (Methods), 1,425 (0.8% of all introns) had non-canonical terminal dinucleotides. Remarkably, 71% (1,014/1,425) of non-canonical introns were classified as either confident (60%) or likely (11%) U12-type introns (Fig. 2A, S7A). These non-canonical U12-type introns were dominated by boundary pairs with a single difference from canonical pairs, in particular AT-AG (29%), AT-AA (27%), GT-AT (17%), and AT-AT (8%). As with the canonical U12-type introns above, the intronic and U12-type character of these introns was supported by conservation across *P. polycephalum* paralogs (Methods, Figs. S7B-C).

**Fig. 2.**
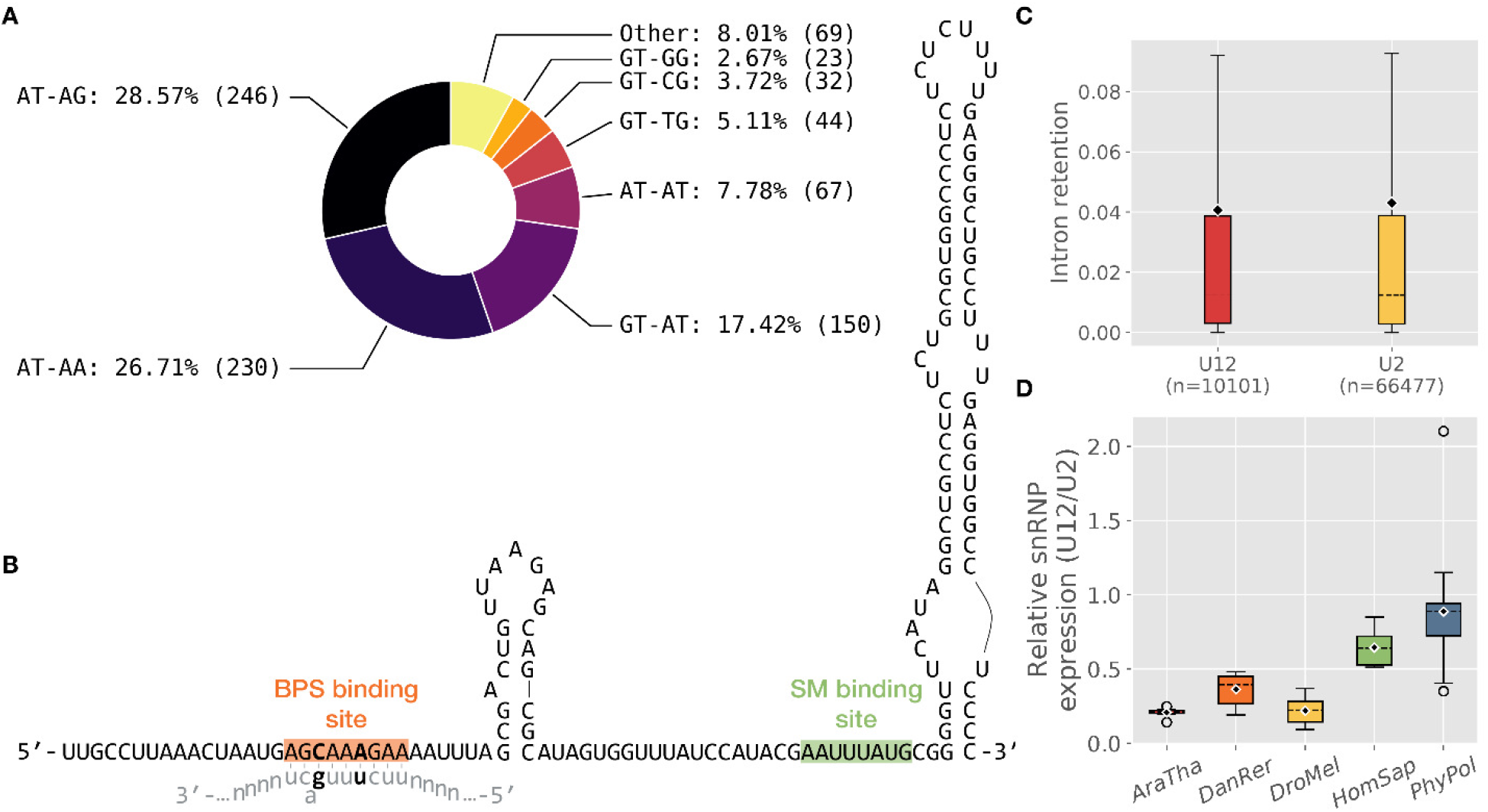
Transformed features of the *P. polycephalum* minor splicing system. **(A)** Non-canonical U12-type intron splice boundaries. **(B)** Putative U12 snRNA sequence and secondary structure (structure based on [15]). Highlighted are the BPS binding site (orange) and the SM binding site (green) and the consensus intronic branchpoint motif (lowercase). The BPS binding site contains two changes relative to the canonical U12 snRNA (bold) which exactly complement changes in the putative TTTGA BPS motif relative to the canonical motif (also bold). **(C)** Comparison of intron retention in RNA-seq data for U12-type and U2-type introns. In contrast to mammals (Figure S7), average intron retention of U12-type introns is not higher than that of U2-type introns in *P. polycephalum*. **(D)** Increased expression of the U12 spliceosome in *P. polycephalum*. The average expression of U12 spliceosomal components, relative to U2 spliceosomal components, is significantly higher in *P. polycephalum* than other species (Methods). For both **C** and **D**, dashed line = median, diamond = mean, whiskers = 1.5 IQR.

We also scrutinized components of the U12 spliceosome in *P. polycephalum*. A genomic search revealed a single candidate for the U12 snRNA that basepairs with the branchpoint (as previously reported in [15]). Strikingly, this sequence exhibits two transition mutations relative to the core branchpoint binding motif (underlined): G<u>C</u>AA<u>A</u>GAA, which produce basepairing potential with the putative <u>T</u>TT<u>G</u>A branchpoint with a bulged A, comparable to the canonical structure (Fig. 2B). This apparent coevolution of core U12 spliceosomal machinery and branchpoint sequence represents to our knowledge the first true instance of coevolution of core intronic splicing motifs and core spliceosomal snRNAs.

U12-type introns in other species have been reported to have lower splicing efficiency than U2-type introns [12,13,47–49], raising the question of how *P. polycephalum* copes with ubiquitous U12-type introns. To investigate, we used RNA-seq and IRFinder [40] to calculate intron retention, and estimated splicing efficiency by comparing fractions of spliced and unspliced junction support between U2- and U12-type introns (Methods). Surprisingly, U12-type introns show slightly lower average intron retention (and higher average splicing efficiency) when compared either *en masse* (Figs. 2C, S8) or in matched pairwise comparisons with neighboring U2-type introns in the same gene (Figs. S9-10). These data suggest that minor spliceosomal kinetics are not inherently less efficient, and that they have been optimized in *P. polycephalum* in concert with the spread of U12-type introns. Consistent with increased efficiency of the U12 machinery in this lineage, we also found that the difference in average expression between the U12 and U2 spliceosomal components was smaller in *P. polycephalum* than is the case in species with lower U12-type intron densities (Fig. 2D).

### U12-type intron creation in *Physarum*

The near absence of U12-type intron creation in most lineages has been argued to reflect the low likelihood of random appearance of the strict U12-type splicing motifs at a given locus [10,50]. How, then, did *P. polycephalum* acquire so many U12-type introns? Inspired by cases of U2-type intron creation by insertion of DNA transposable elements [51,52], we scrutinized U12 splice sites in *P. polycephalum*. We noted that many *P. polycephalum* U12-type introns carry sequences that resemble the signature of DNA transposable elements, namely inverted repeats (rta<u>tctt</u>…<u>aagA</u>TAT) flanked by a direct repeat of a TA motif. This suggests the possibility that *P. polycephalum* U12-type introns could have been created by a novel DNA transposable element with TCTT-AAGA termini and TA insertion site (Fig. 3). It is of note that *P. polycephalum* U12-type introns differ at two sites from the corresponding classic motif (TC<u>C</u>T-<u>Y</u>AGA), where both changes increase the repeat character. An ancestral decrease in the length and stringency of the branchpoint motif could have increased the probability of *de novo* evolution of a DNA transposable element carrying sufficiently U12-like splice sites for new insertions to be recognized by the U12 spliceosome.

**Fig. 3.**
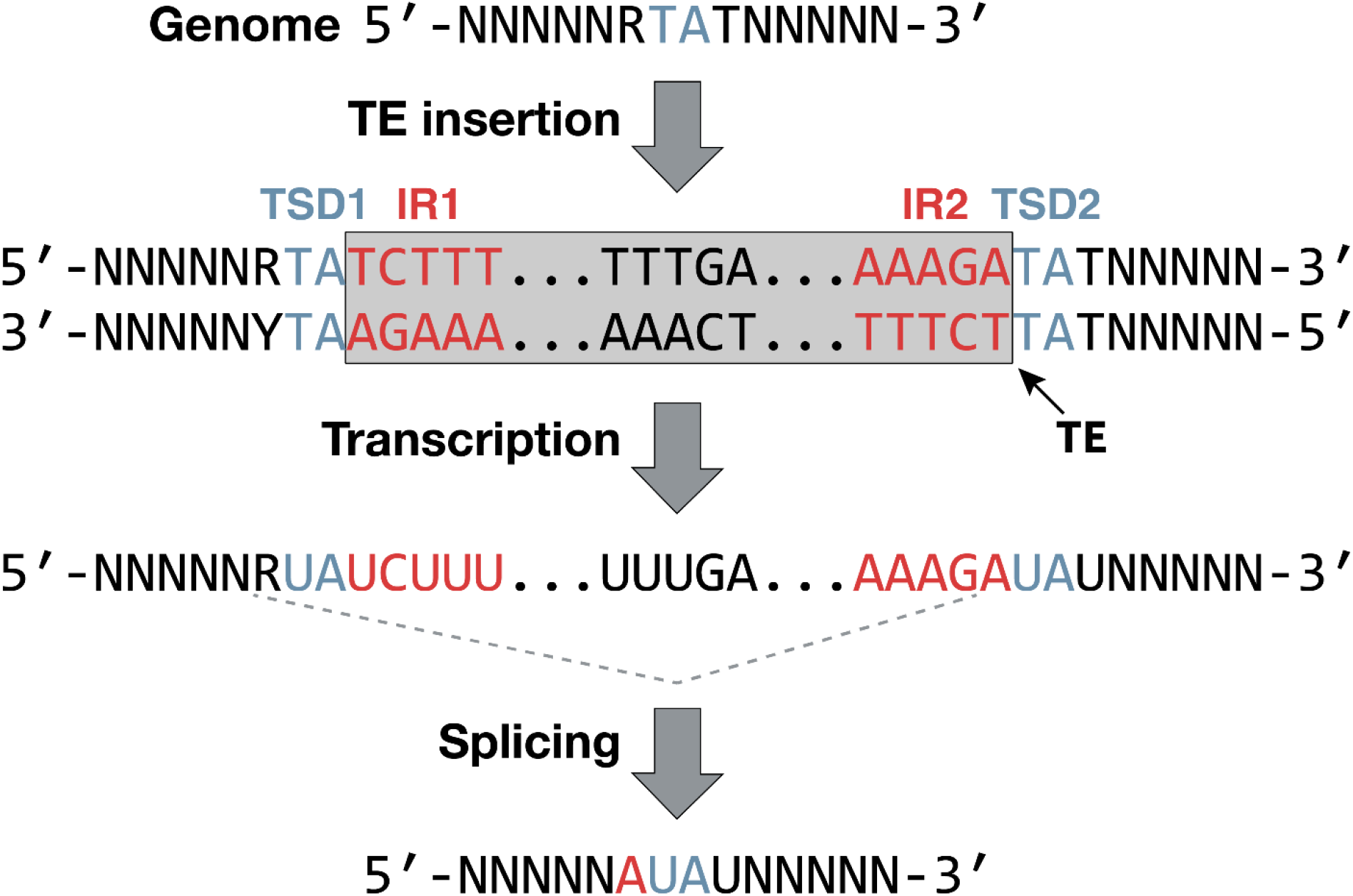
Proposed mechanism for transposon-driven creation of U12-type introns in *P. polycephalum*. Insertion of a transposable element (TE, gray box) carrying inverted repeats (IR1/IR2, red) leads to duplication of a TA target side (TSD1/TSD2, blue). Splicing at RT-AG boundaries leads to a spliced transcript with a sequence identical or nearly identical to the initial gene sequence with loss of an R (G/A) nucleotide and gain of the 3′ A from the transposable element, maintaining the original reading frame.

## Discussion

In contrast to the portrait of the U12 spliceosomal system as rare, ancient, static and suboptimal, these results expand our understanding of U12 diversity, by (i) increasing the upper bound of U12-type intron density per species by two orders of magnitude; (ii) showing that U12-type introns have been gained *en masse* through eukaryotic evolution; and (iii) showing that U12-type splicing is not necessarily less efficient than U2-type splicing. *P. polycephalum* provides promise for an understanding of the flexibility of U12 splicing, a potentially important role given the increasing appreciation of U12 splicing errors in development and human disease. In addition, further study of *P. polycephalum* and related species may provide insights into the mechanisms and functional implications of the initial proliferation of introns in ancestral eukaryotes.

## Author contributions

M.E. supplied initial supporting data, G.E.L. and S.W.R. conceived of the study, G.E.L. performed computational analyses, data processing and visualization, G.E.L. and S.W.R. analyzed and interpreted the results, G.E.L and S.W.R. wrote the manuscript with input from M.E..

## Materials and Correspondence

Scott W. Roy;

## Data availability

The *Physarum polycephalum* genome and annotation file used in our analyses have been uploaded to the following Zenondo archive: https://doi.org/10.5281/zenodo.4086119; intron coordinates and U12-type probability scores for all *P. polycephalum* introns have been archived at https://doi.org/10.5281/zenodo.4099156.

## Code availability

The intron classifier intronIC is open-source and available on GitHub: https://www.github.com/glarue/intronIC; the modified version used in this manuscript for classifying introns in *P. polycephalum* specifically has been archived at https://doi.org/10.5281/zenodo.4265109.

## Supplementary Figures

**Figure S1.**
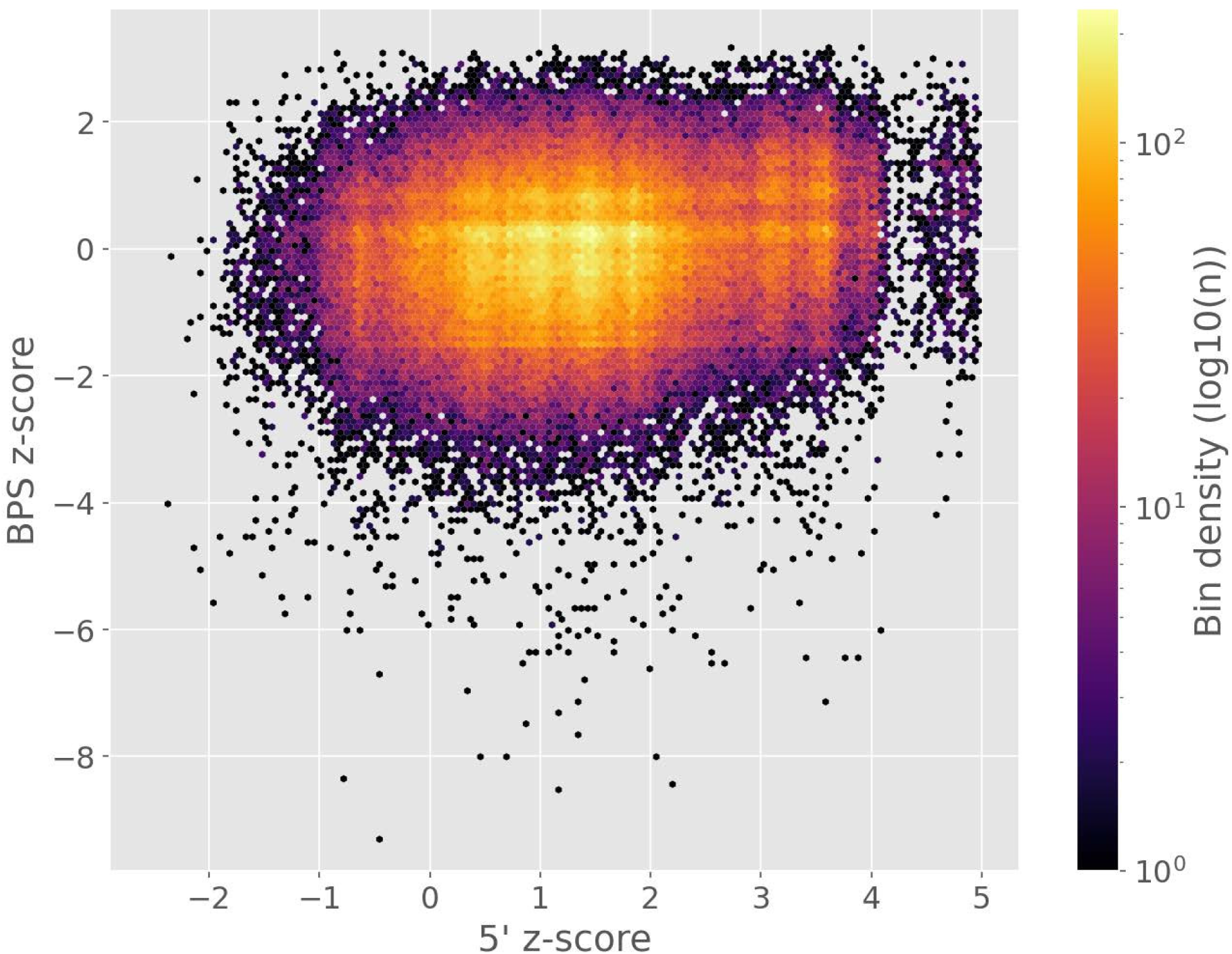
BPS-vs-5′SS score plot for all *P. polycephalum* introns under default settings via intronIC [1]. The default PWMs used by intronIC are derived from human introns, and for divergent motifs like those present in *P. polycephalum* (especially the BPS motif) they fail to produce clear differentiation (i.e. separation of U12-type introns into a distinct cloud in the first quadrant). Curation of species-specific PWMs for *P. polycephalum* resulted in clearer differentiation along both axes (as in Figure 1D).

**Figure S2.**
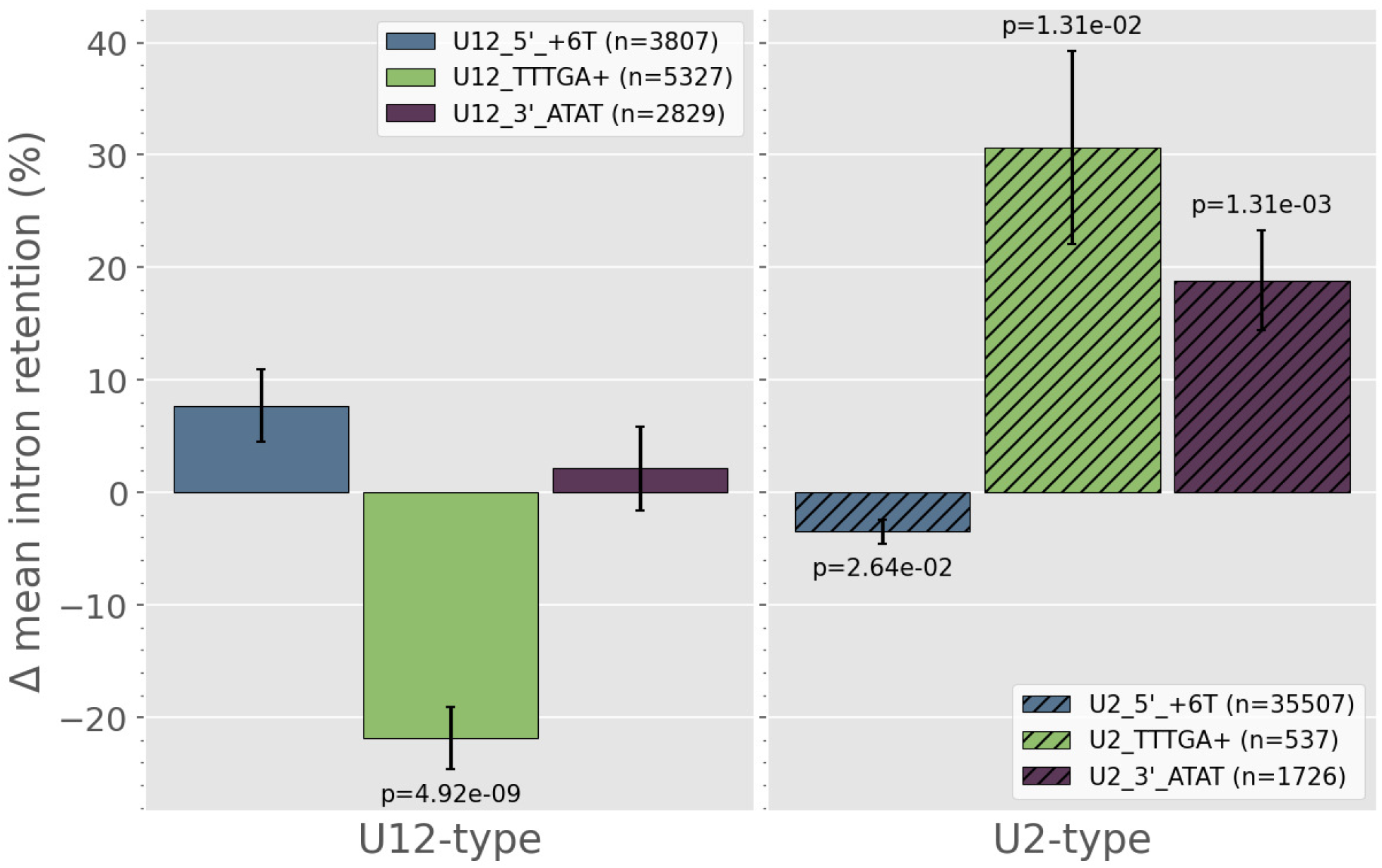
Relative intron retention for U12- (left) and U2-type (right) introns based on sequence features. Differences from the mean for each category are relative to all other introns of the same type. A negative/positive value indicates that introns with the given feature exhibit more/less efficient splicing relative to other introns of the same type. Features shown are “5′_+6T”, introns with a T at position +6 in the intron; “TTTGA+”, introns with the TTTGA motif within the last 55 bases of the intron; “3′_ATAT”, introns with the motif ATAT immediately downstream of the 3′SS.

**Figure S3.**
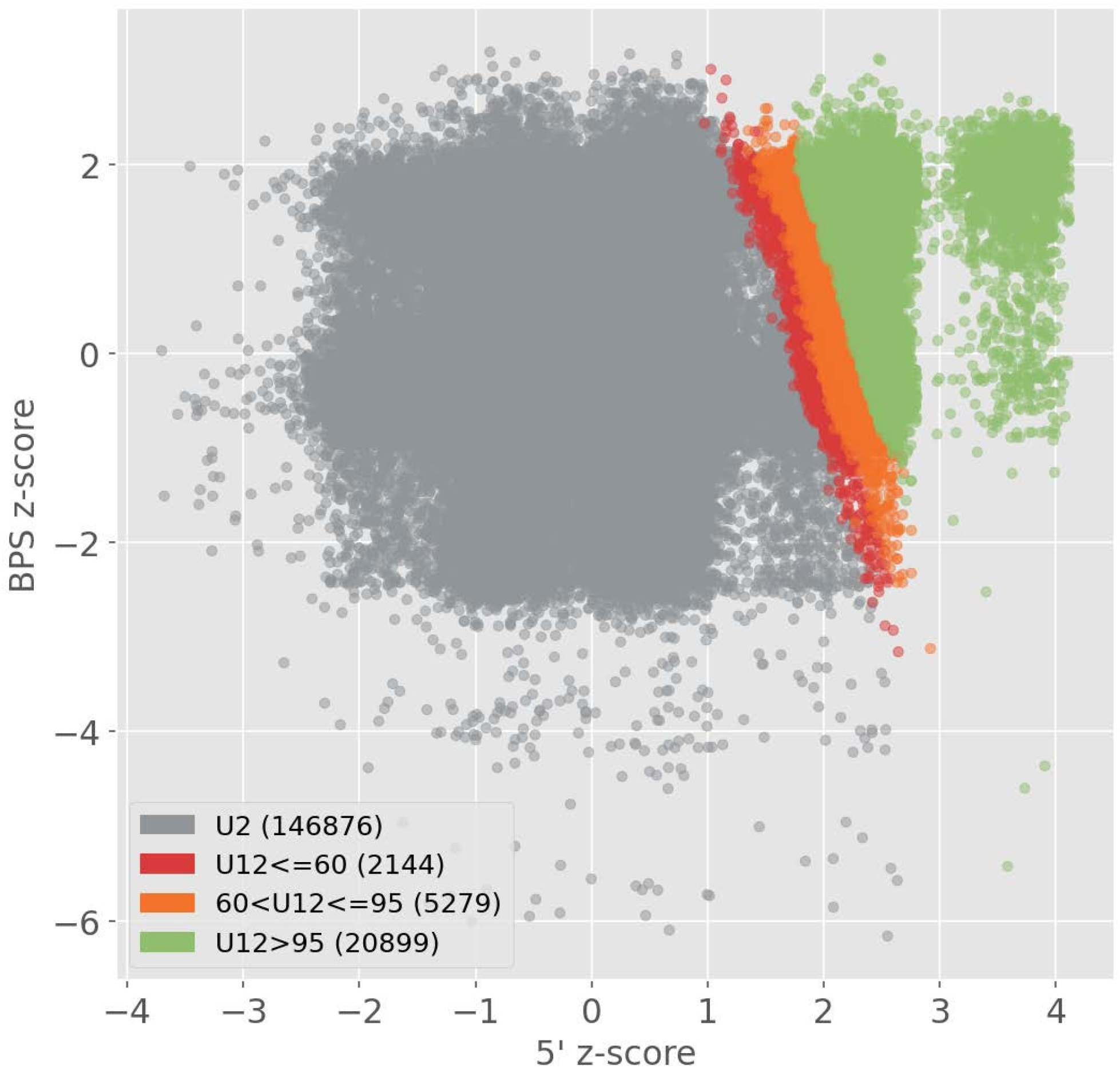
BPS-vs-5′SS score plot with assigned classifications for all *P. polycephalum* introns. The same underlying data as Figure 1C, where each point represents an intron, and the color indicates the U12-type probability classification (gray, U2-type; red, U12-type with probability ≤ 60%; orange, U12-type with probability 60-95%; green, U12-type with probability > 95%). Throughout our analyses, only the > 95% category were considered U12-type (unless explicitly stated otherwise).

**Figure S4.**
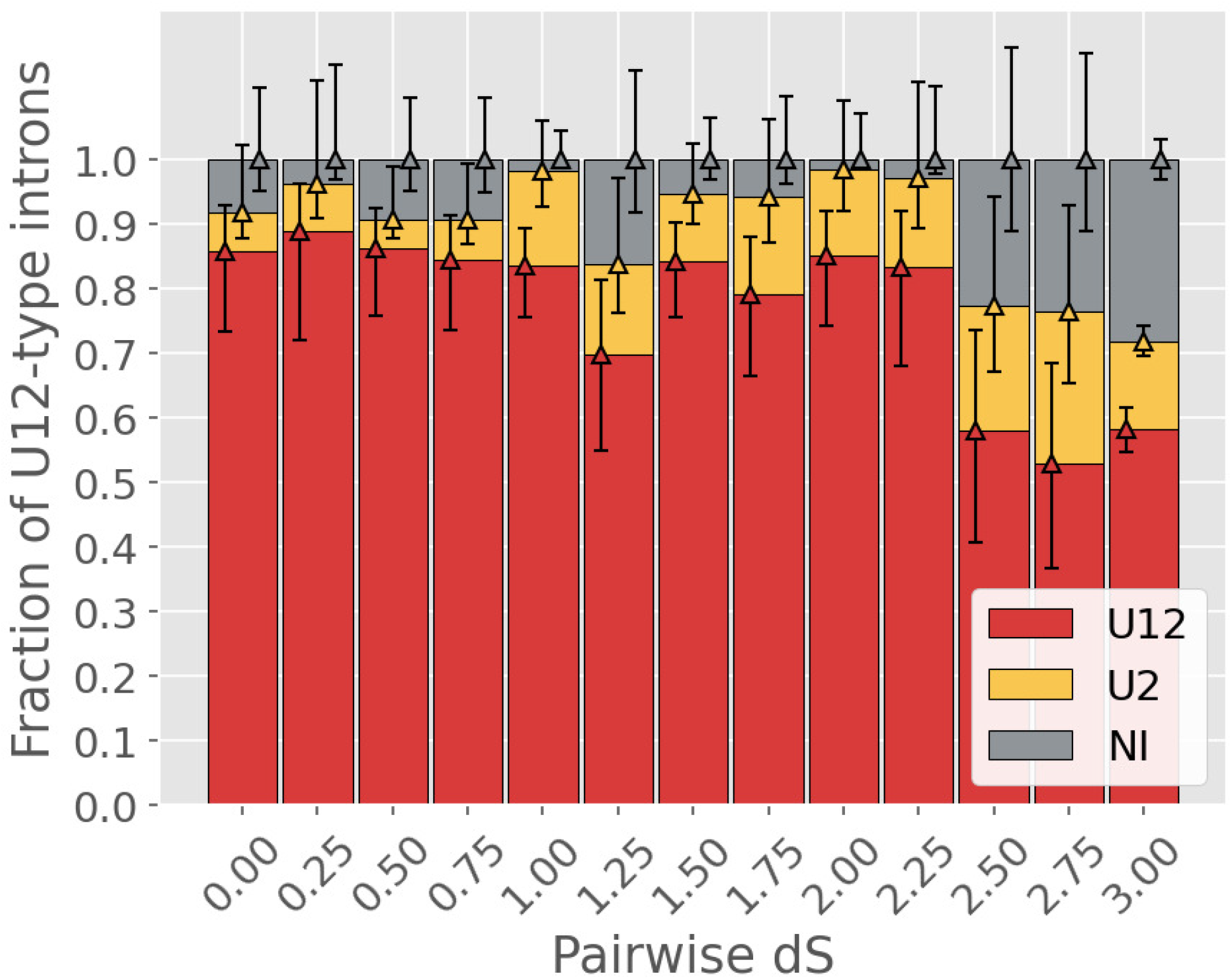
Between-paralog comparison provides little evidence for ongoing U12-type intron gain in *P. polycephalum*. For U12-type intron-containing paralog pairs sharing at least one intron of either type (to exclude recent retrogenes), pairwise dS values were used to bin all pairs into the range [0, 3]; dS values >=3 were binned together. Within a given bin, each U12-type intron has one of three possible conservation states in its corresponding paralog: U12-type (red), U2-type (yellow) or no intron present (“no intron”, gray). These data suggest that there have not been major U12-type intron gains in *P. polycephalum* since a time corresponding to at least dS ∼2.5. Whiskers represent the binomial proportion confidence intervals (Wilson score intervals) for the three categories (category indicated by color of associated diamond).

**Figure S5.**
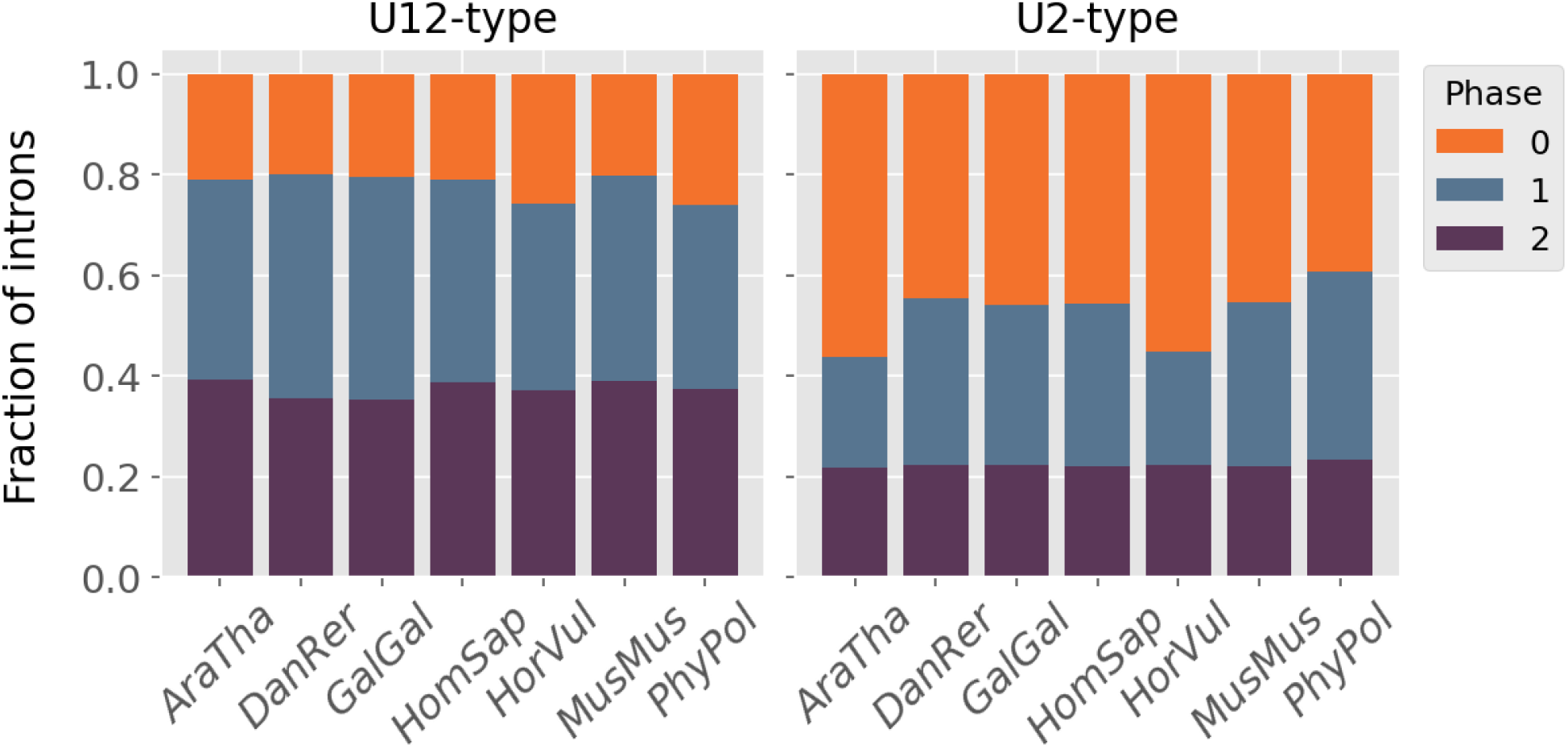
Phase distribution of U12- (left) and U2-type (right) introns across different species. U12-type introns in *P. polycephalum* (*PhyPol*) display a bias away from phase 0, as in other species, whereas U2-type introns show a bias against phase 2. For each species, only introns interrupting coding sequence from the longest isoform of each gene were included. See Table S1 for species abbreviations.

**Figure S6.**
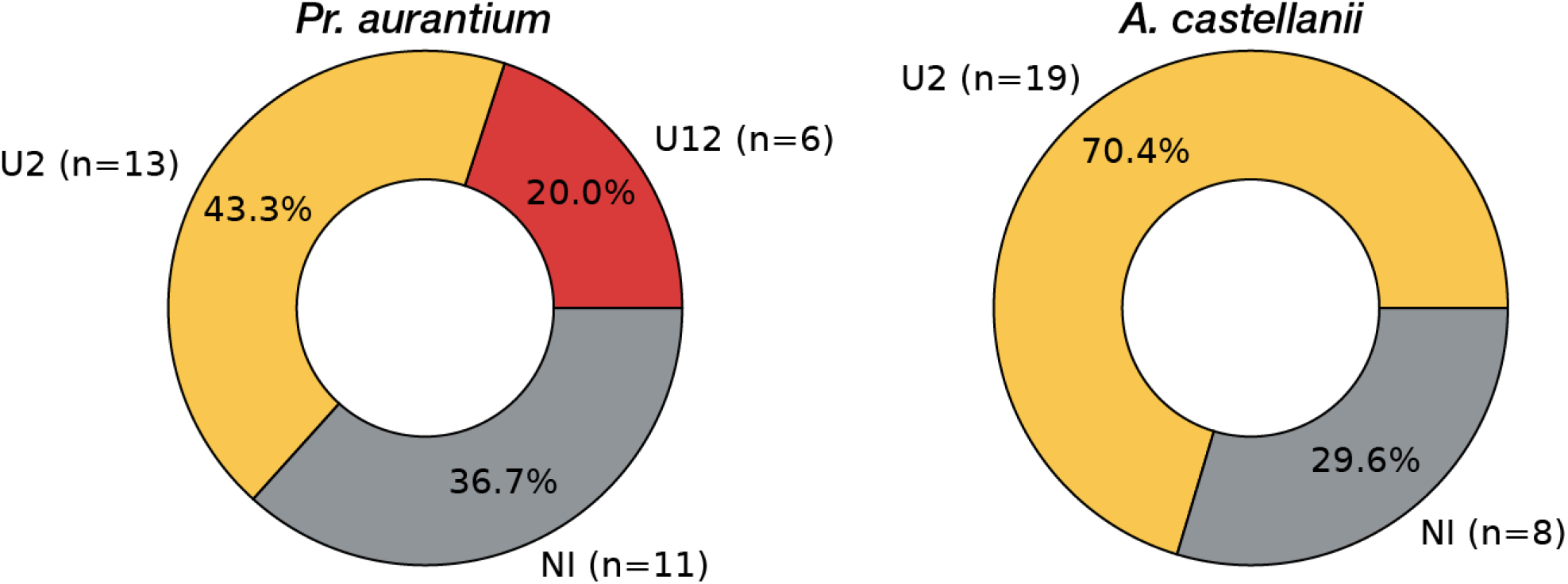
Ancestral U12-type introns in *P. polycephalum* are conserved as introns in other amoebozoans. Each pie chart shows the conservation status (red, U12-type; yellow, U2-type; gray, no intron) of the same ancestral set of *P. polycephalum* U12-type introns (introns conserved as U12-type with one or more non-amoebozoans) in the variosean amoeba *Protostelium aurantium* (left) and the discosean amoeba *Acanthamoeba castellanii* (right). In each case, a significant majority of the U12-type introns are conserved as introns. This data suggest that these species have not undergone massive loss of U12-type introns, and thus that the unprecedented number of U12-type introns in *P. polycephalum* represents significant U12-type intron creation in *P. polycephalum* rather than loss in related species.

**Figure S7.**
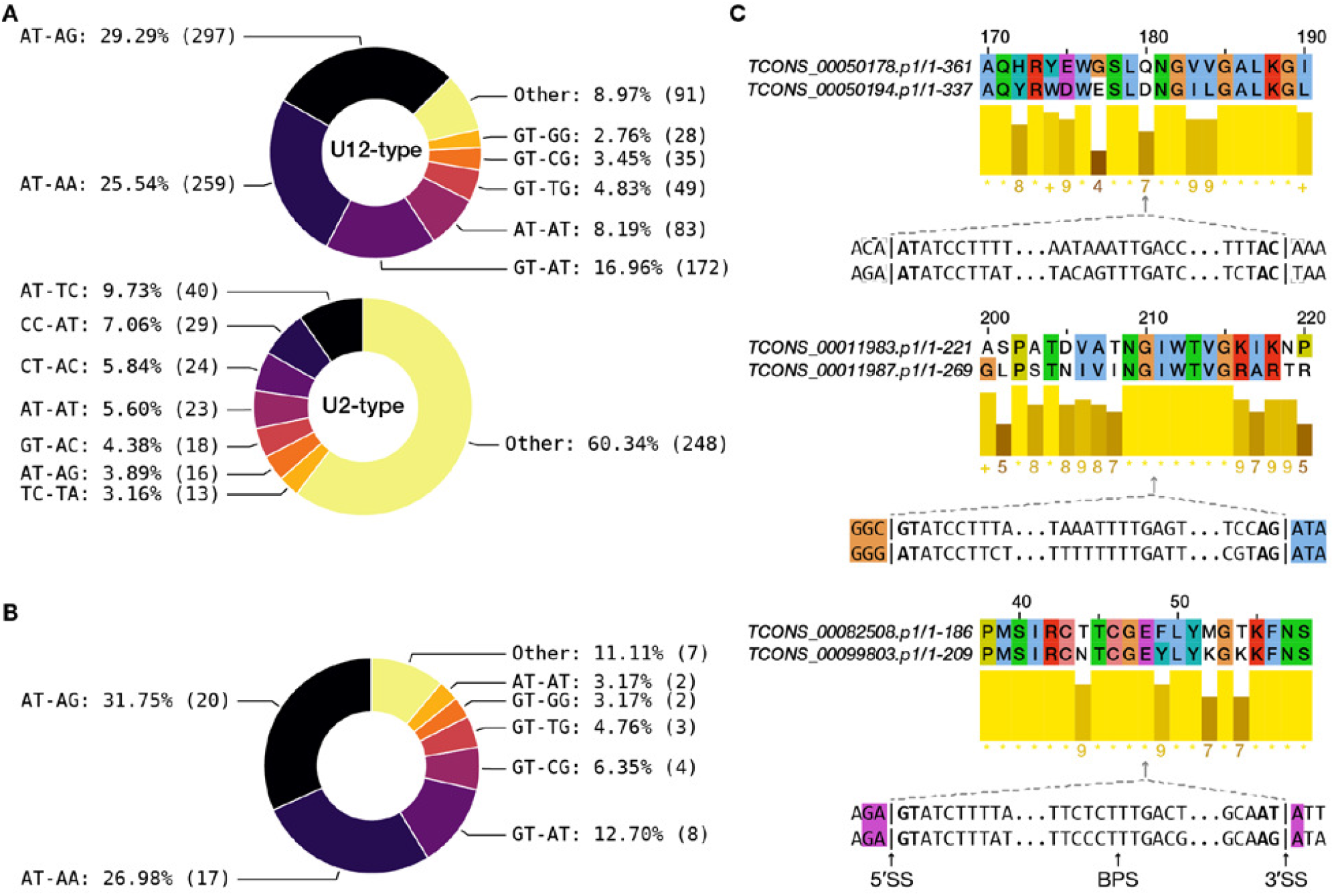
Non-canonical introns in *P. polycephalum*. **(A)** U12- (top) and U2-type (bottom) non-canonical intron subtypes (using a 60% probability threshold for the U12/U2-type classification instead of the 95% threshold used elsewhere e.g. Figure 2A, thereby including “likely” U12-type introns), highlighting the degree to which non-canonical U12-type introns are greatly enriched for a subset of boundary pairs. By contrast, the U2-type non-canonical subtype distribution is much more diffuse. **(B)** Distribution of the subset of non-canonical U12-type introns which are found in regions of good alignment between pairs of *P. polycephalum* paralogs (but not necessarily conserved as introns between pairs), increasing confidence that they are truly introns, showing general consistency with part A. **(C)** Example alignments of *P. polycephalum* paralogs, showing conserved U12-type introns (canonical and non-canonical). Coloring is based on chemical properties of the amino acids, and bars underneath each alignment represent chemical similarities of the aligned amino acids. Colored nucleotides before and after the intron splice sites correspond to the colors of the amino acid(s) in the alignment that are interrupted by the shared intron position. Transcript names appear in italics.

**Figure S8.**
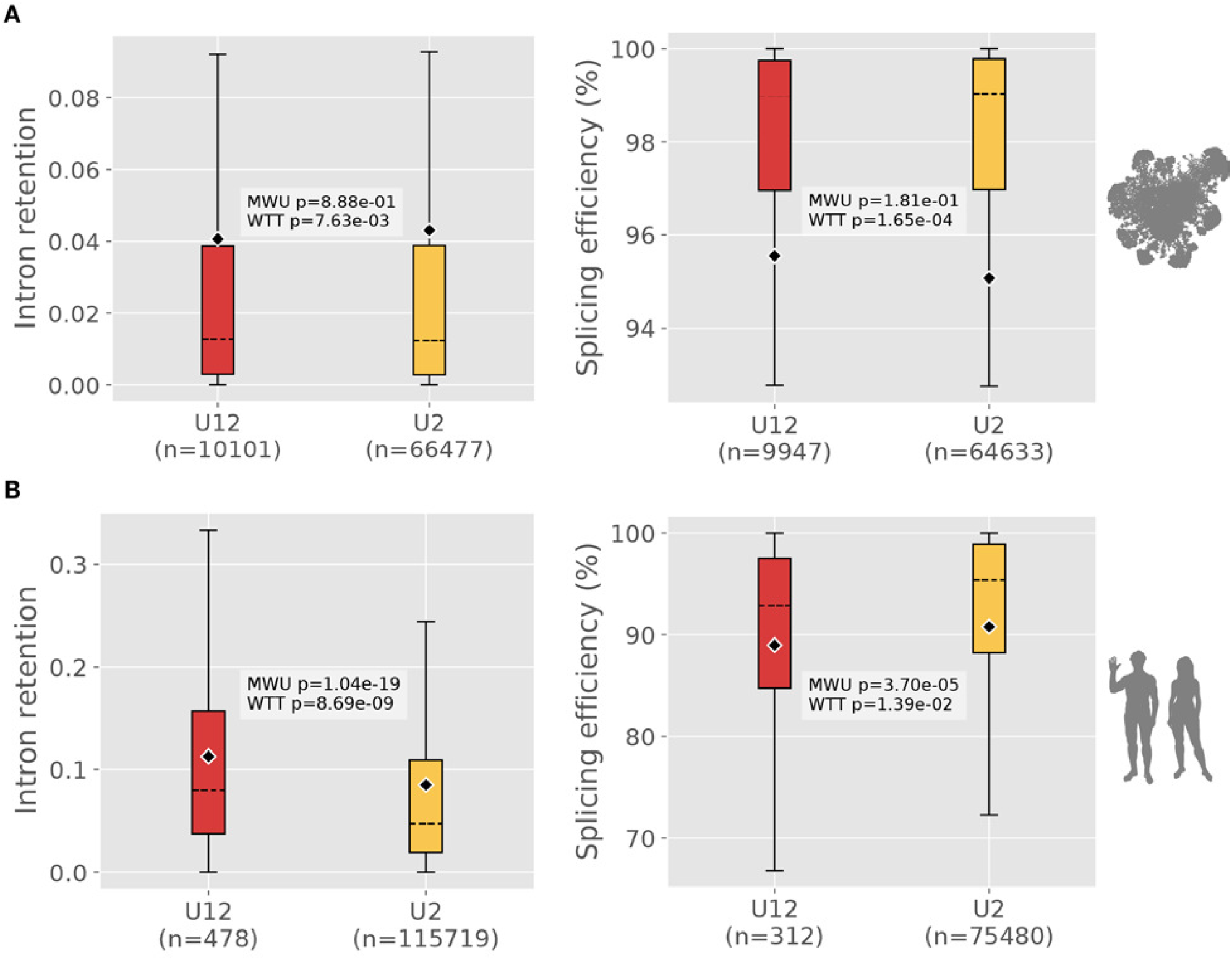
Comparison of intron retention (left) and splicing efficiency (right) in *P. polycephalum* and human. **(A)** Box plot of average intron retention and splicing efficiency data for *P. polycephalum* introns, showing that U12-type introns are neither more retained nor less efficiently-spliced than U2-type introns. Note that although the differences in means between U12- and U2-type introns are significant, this difference is inverted relative to data from other species. The left panel is the same as Figure 2C. **(B)** As in (A), but for *Homo sapiens*. Here, by both statistical measures there are significant differences between the two types of introns, with U12-type introns being more retained and less-efficiently spliced as has been reported elsewhere. MWU: two-tailed Mann-Whitney U test; WTT: two-tailed Welch’s t-test. Note that y-axis scales differ between plots. Dashed line = median, diamond = mean, whiskers = 1.5 IQR.

**Figure S9.**
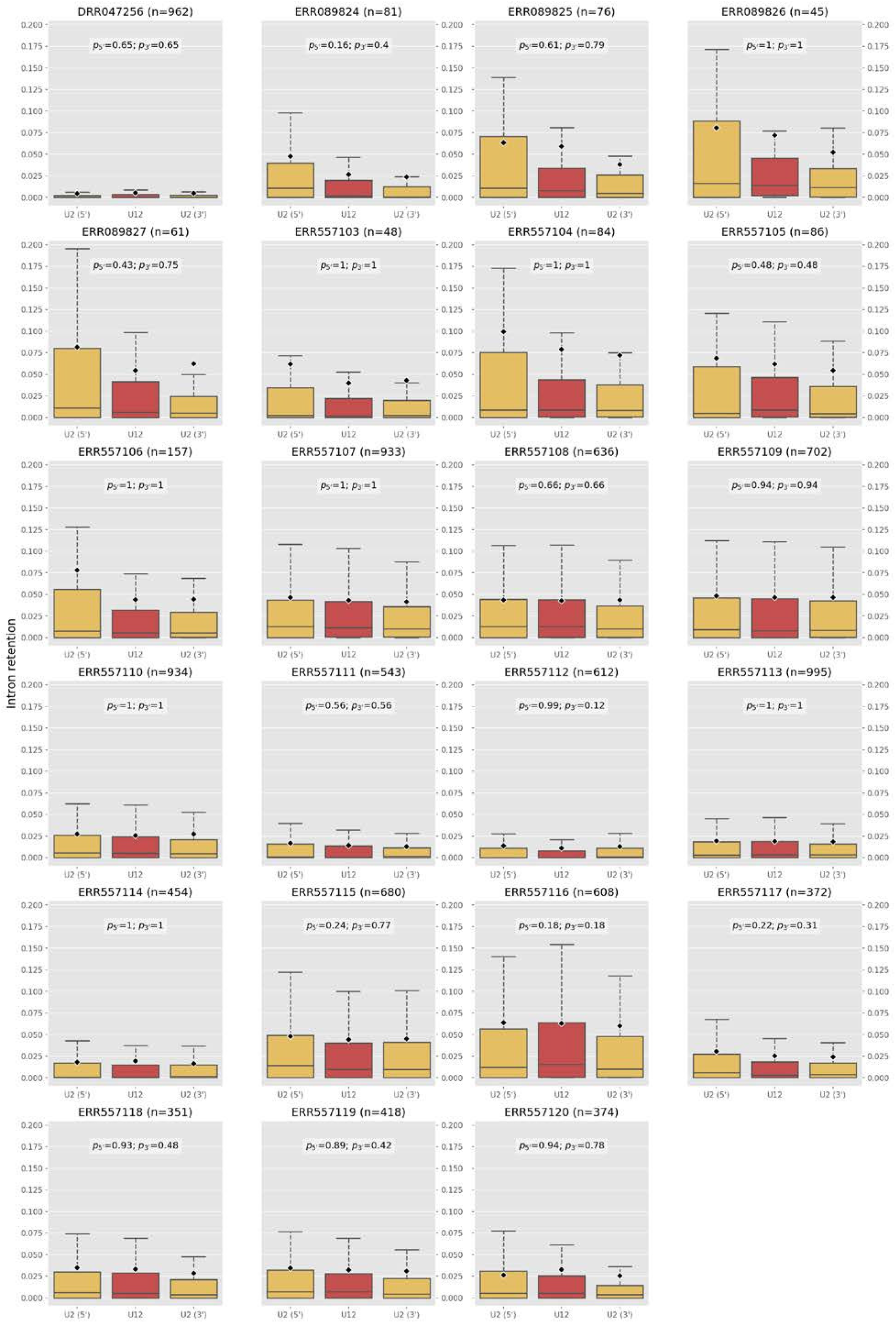
U12-type intron retention is not significantly different from that of neighboring U2-type introns in *P. polycephalum*. Each subplot represents data from a different RNA-seq sample (accession number in subplot title, along with number of introns represented), showing the distribution of intron retention values for U12-type (red) and neighboring U2-type (yellow) introns on both sides (left: 5′, right: 3′). For each neighboring U2-type (defined as introns with U12-type probability scores < 5%) dataset, *p*-values vs. the corresponding U12-type data were obtained via Mann-Whitney U tests, and corrected for multiple testing using the Holm step-down method (reported as *p*5′ and *p*3′ for the 5′ and 3′ U2-type introns, respectively). Dashed line = median, diamond = mean, whiskers = 1.5 IQR.

**Figure S10.**
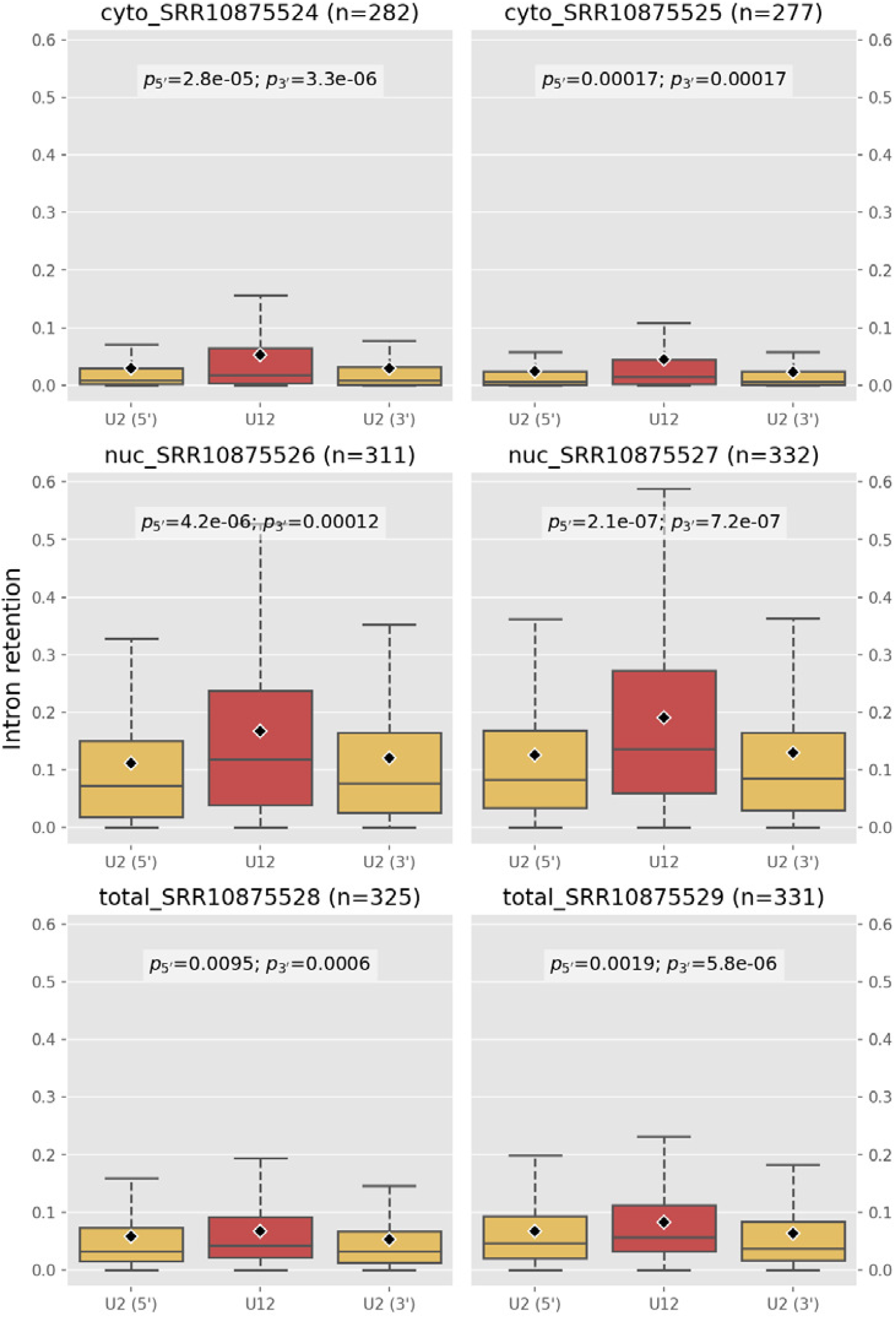
U12-type intron retention is significantly higher than that of neighboring U2-type introns in *Homo sapiens*. RNA-seq sets from three different protocols (cytosolic (“cyto”), nuclear (“nuc”) and total RNA, as indicated in the titles of each subplot) were used to compare U12-type intron retention to that of neighboring U2-type introns. In each case, U12-type introns were more retained than their neighboring U2-type introns, in contrast to our results for *P. polycephalum* (Figure S9). Dashed line = median, diamond = mean, whiskers = 1.5 IQR.

